## Supplemental File 1 for "De novo design of protein minibinder agonists of TLR3"

#### Minibinder 1

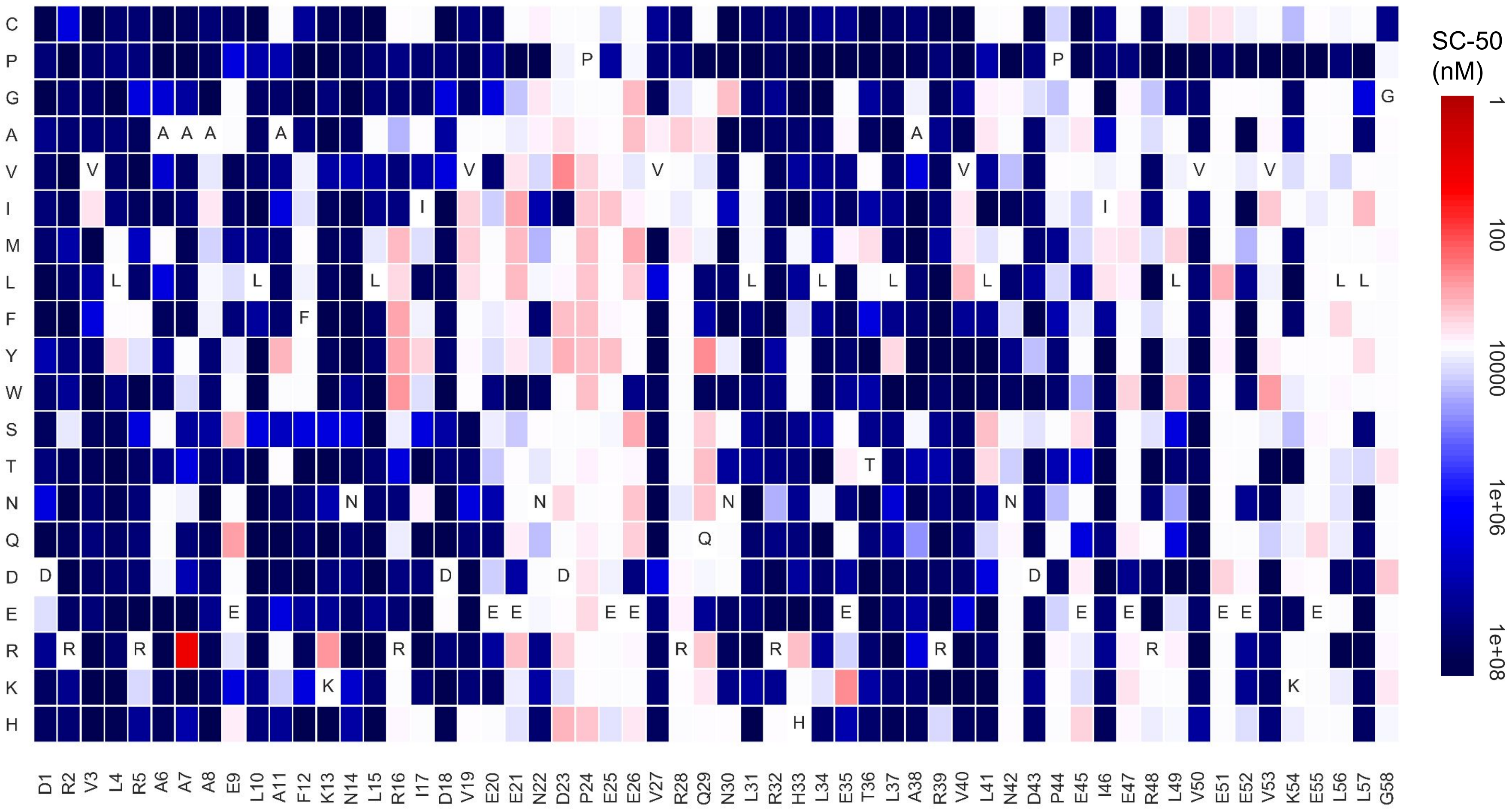

### Minibinder 2

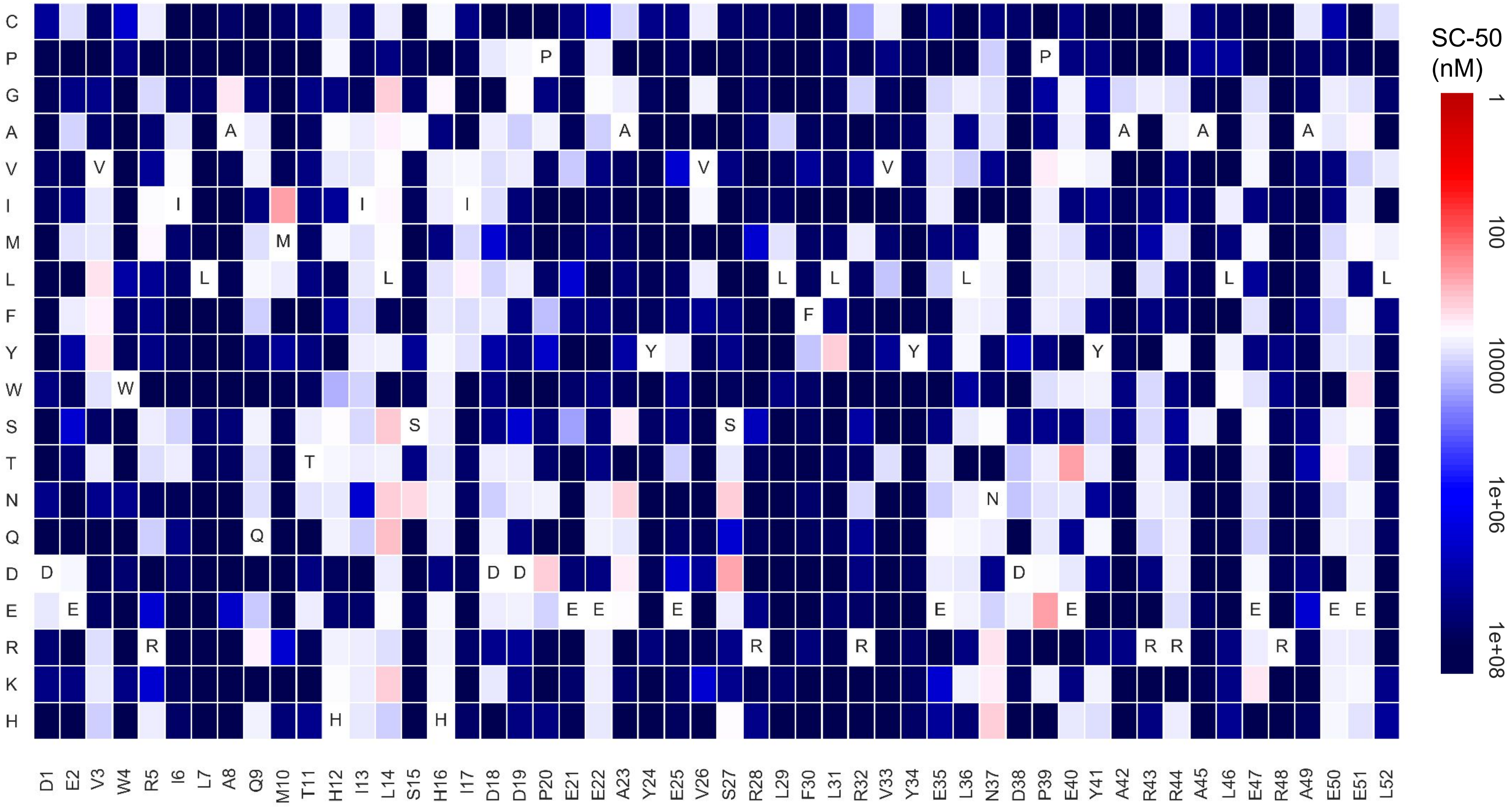

### Minibinder 3

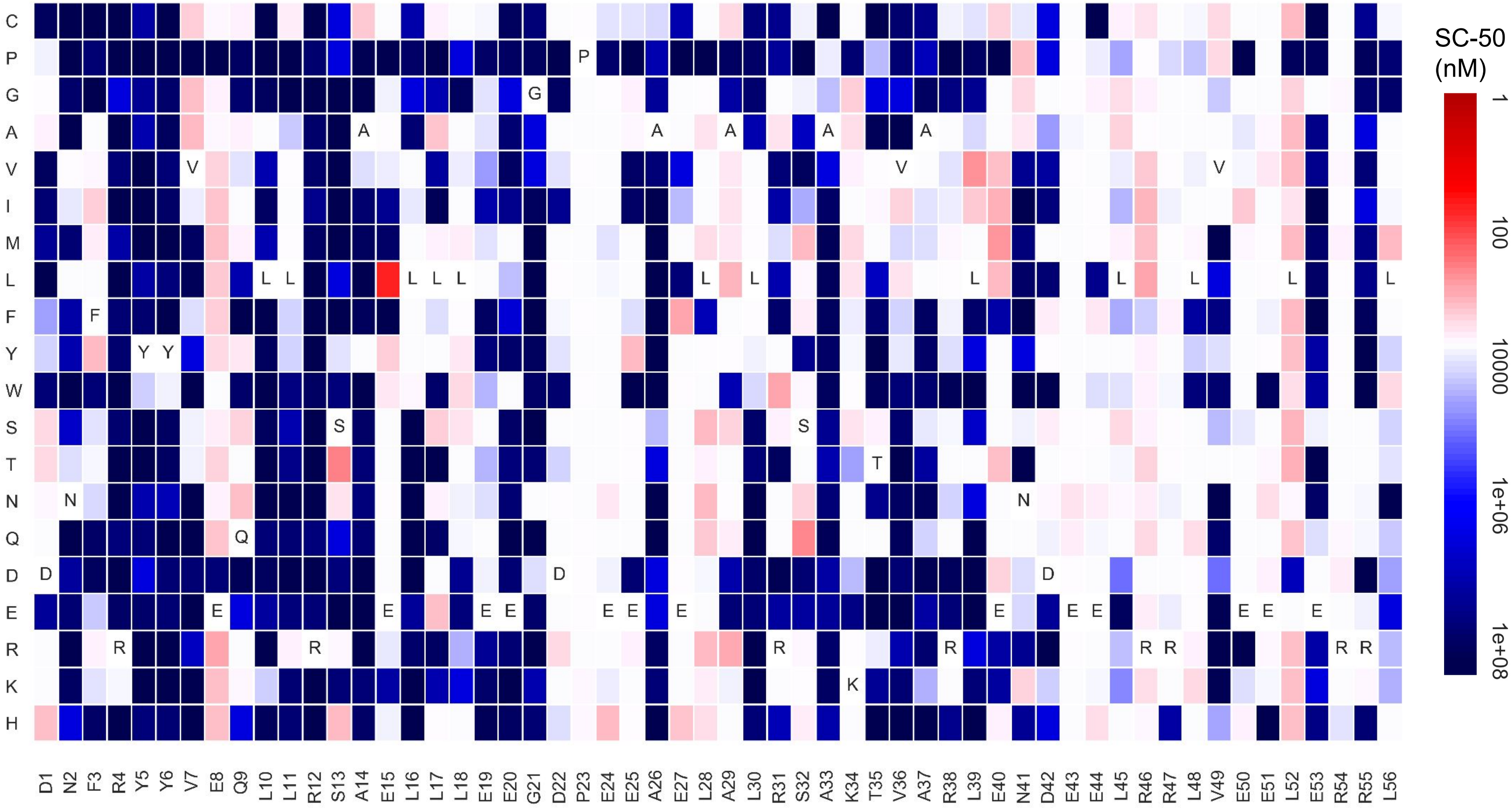

### Minibinder 4

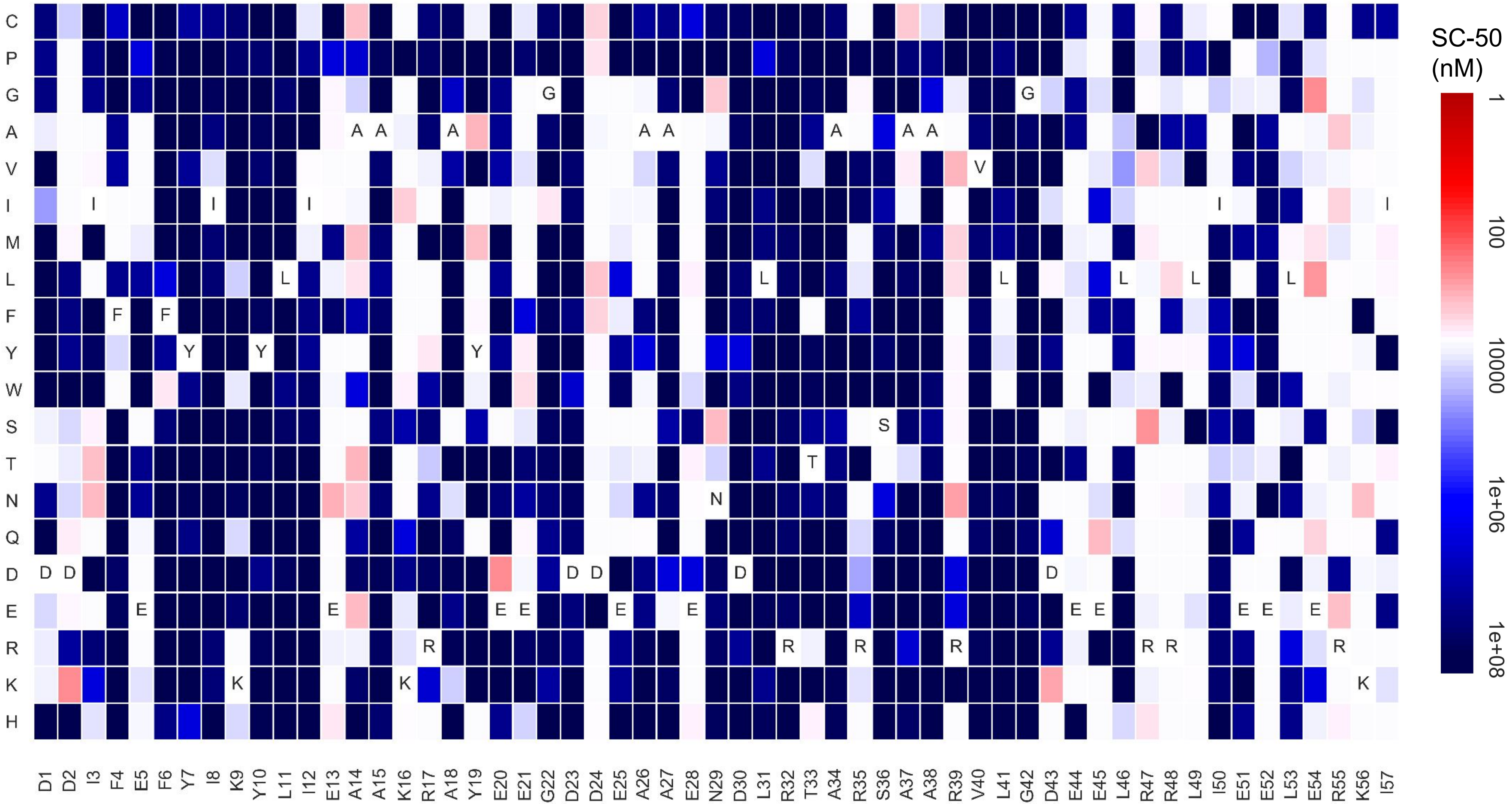

### Minibinder 5

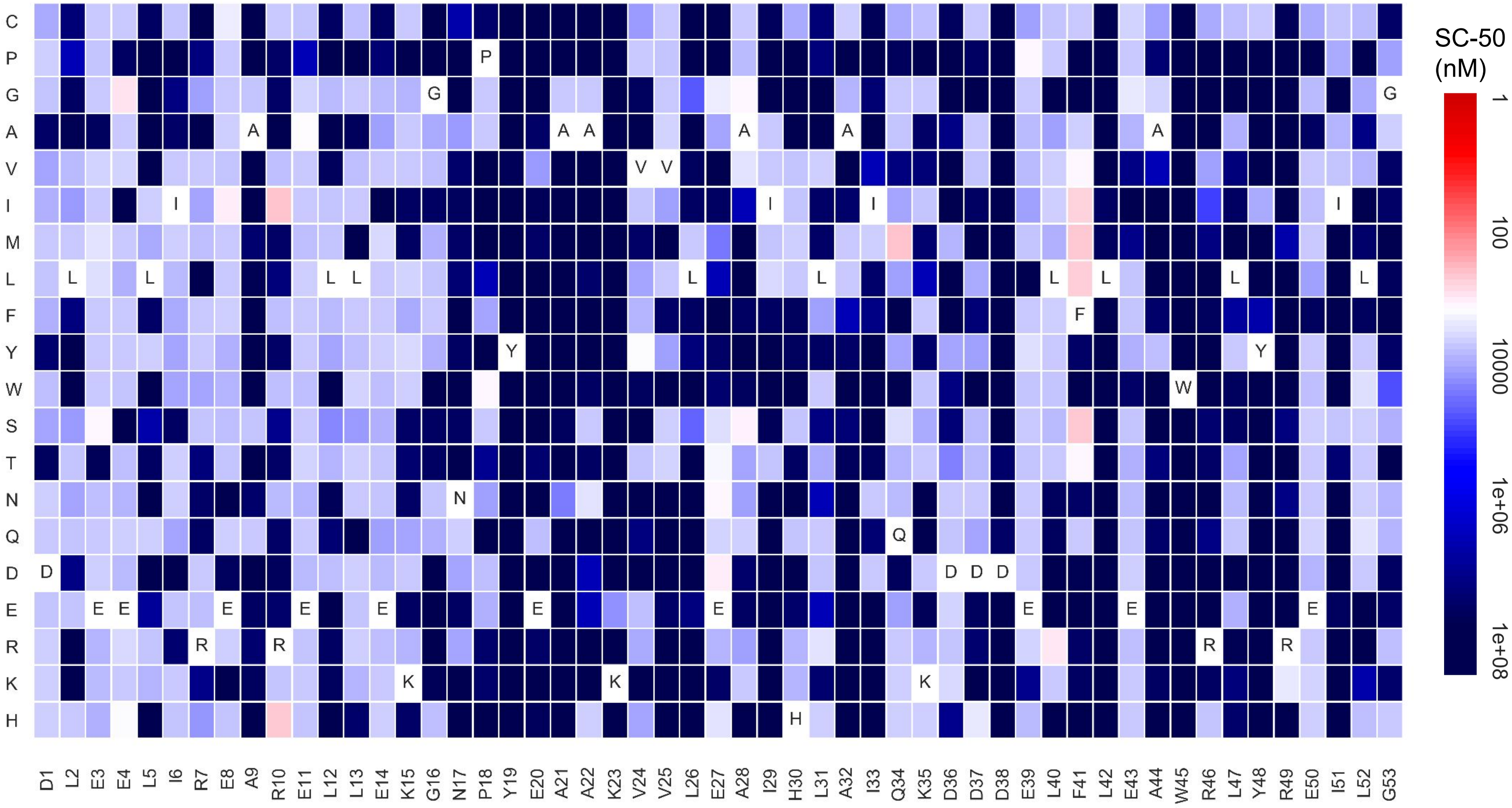

### Minibinder 6

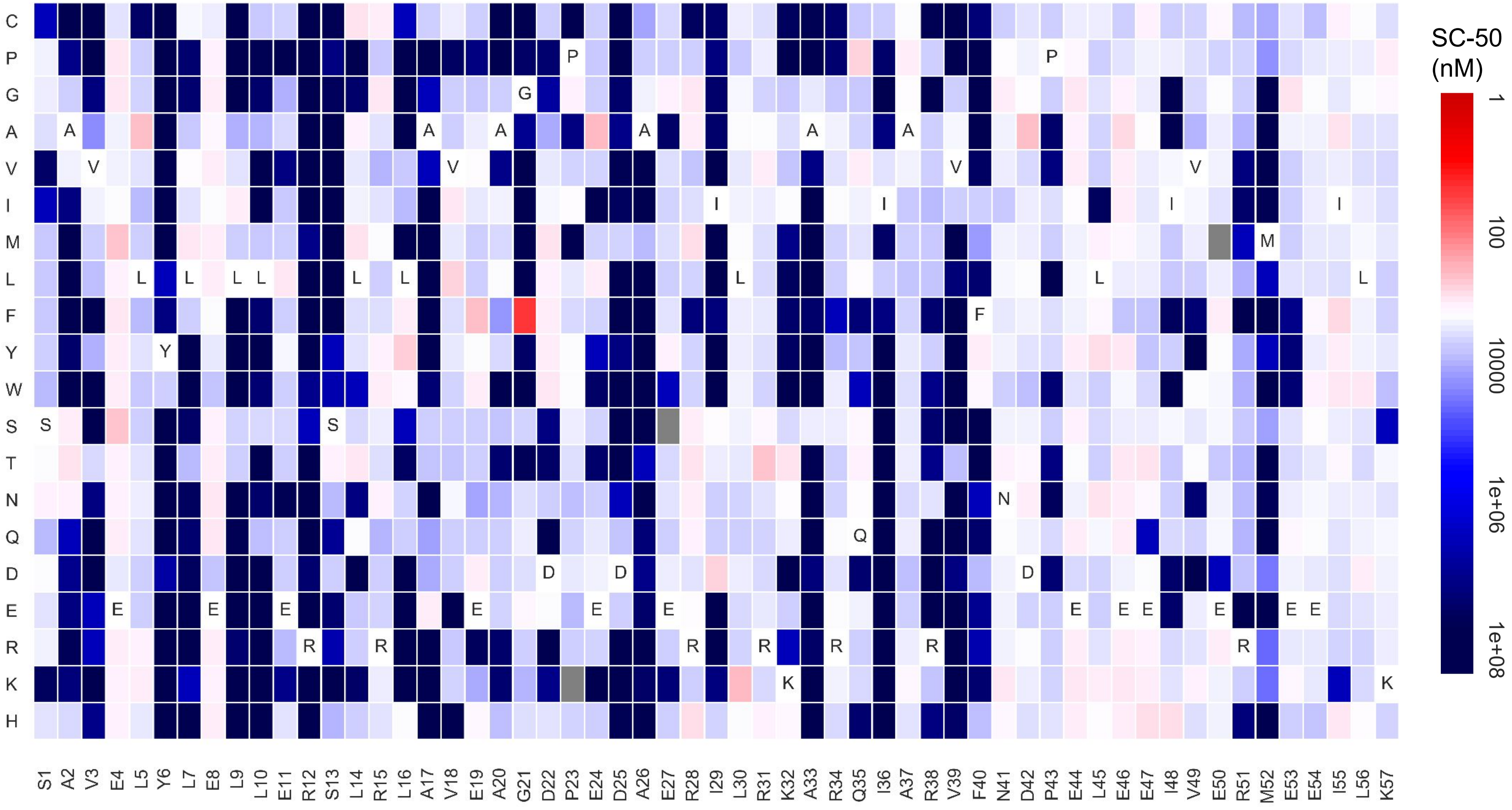

### Minibinder 7

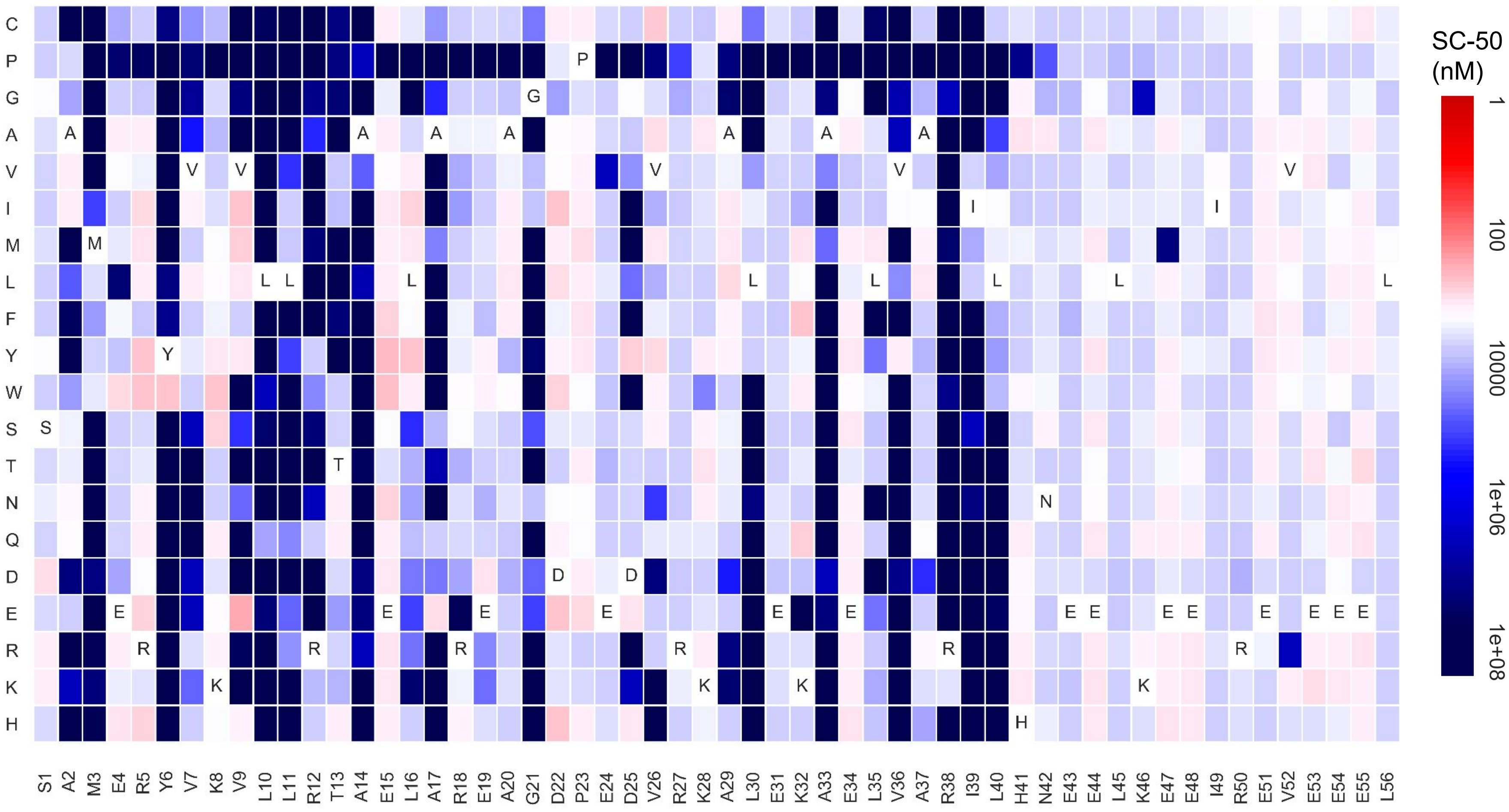

### Minibinder 8

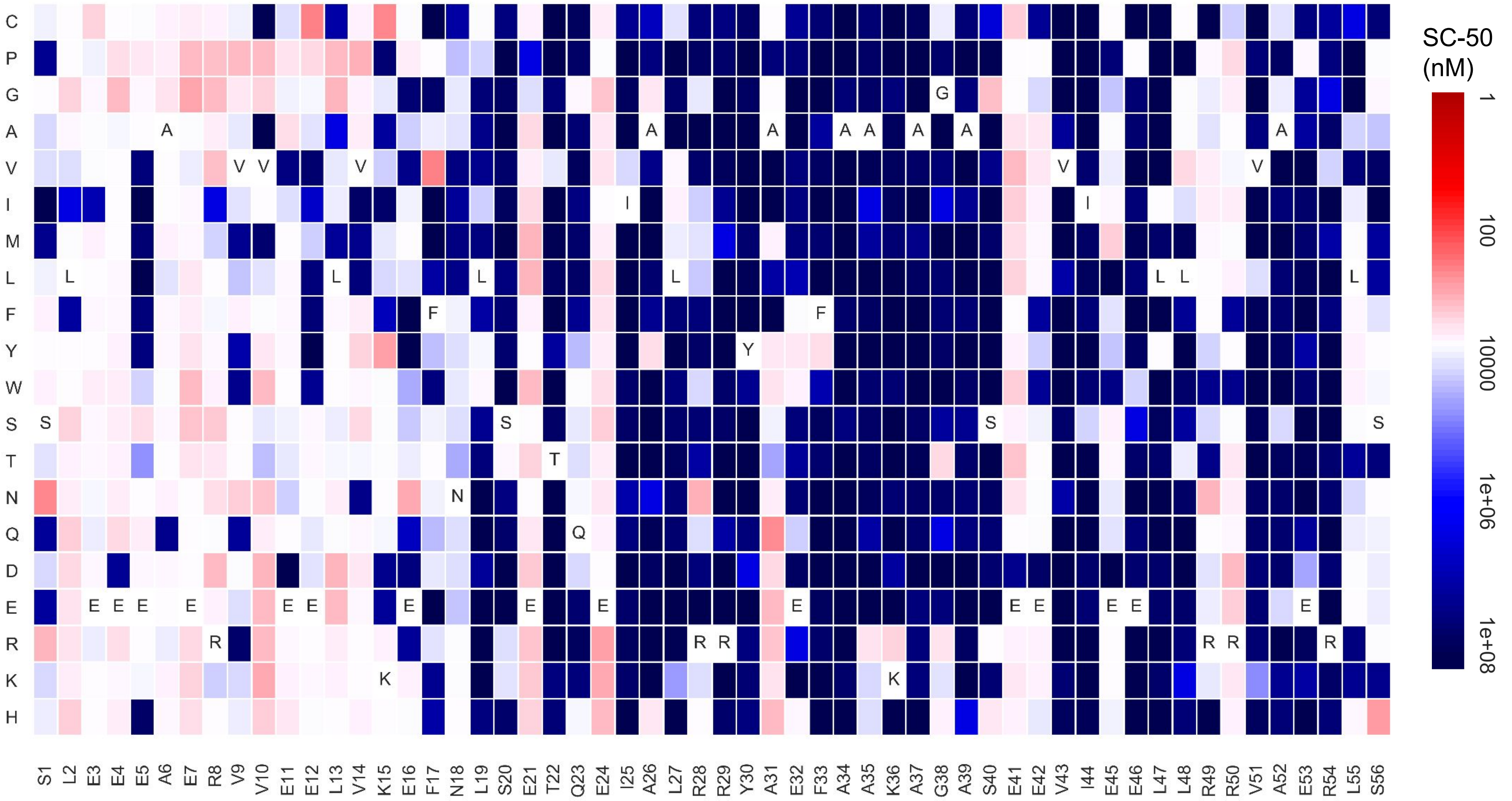

### Minibinder 9

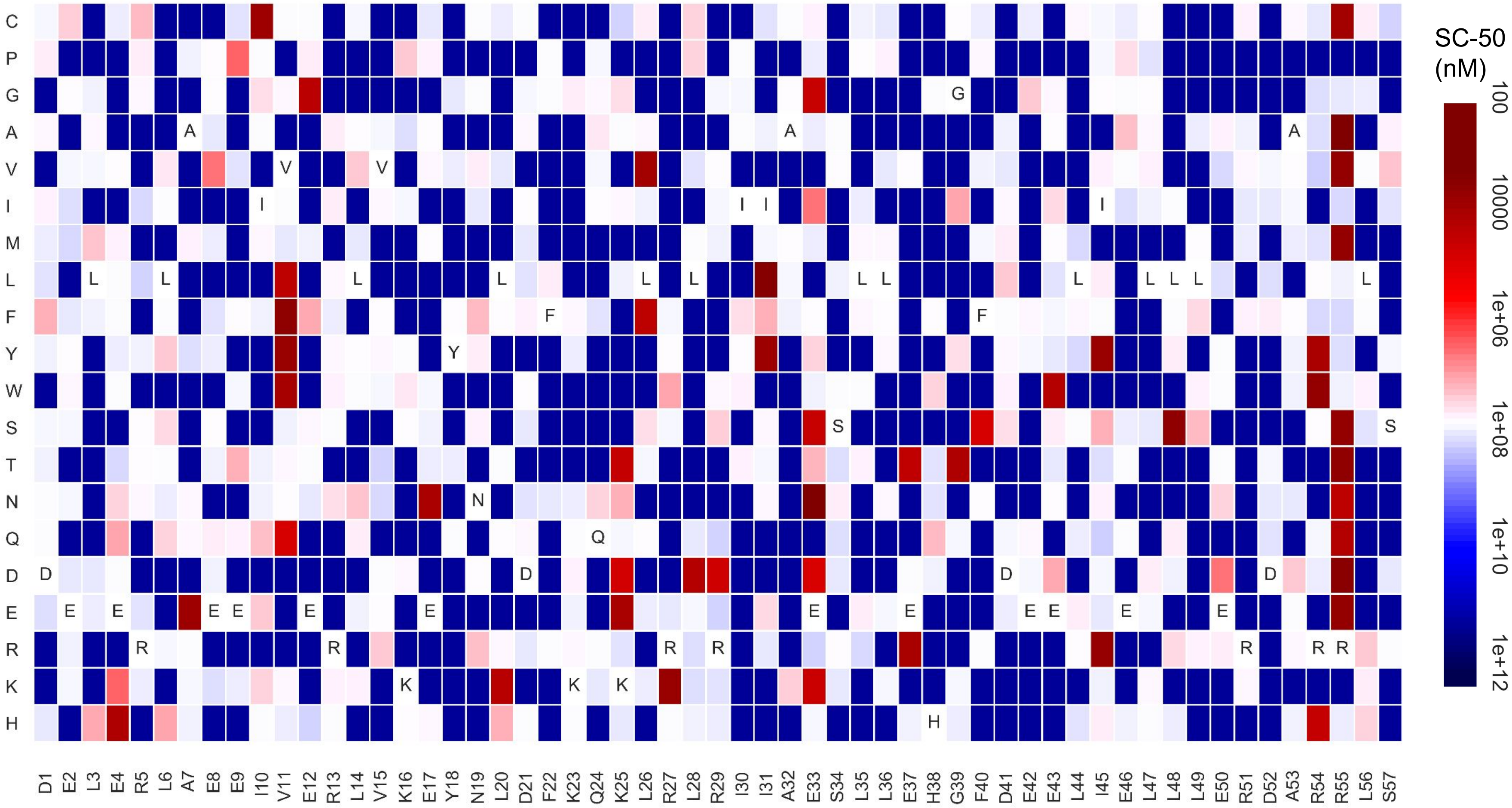

### Minibinder 10

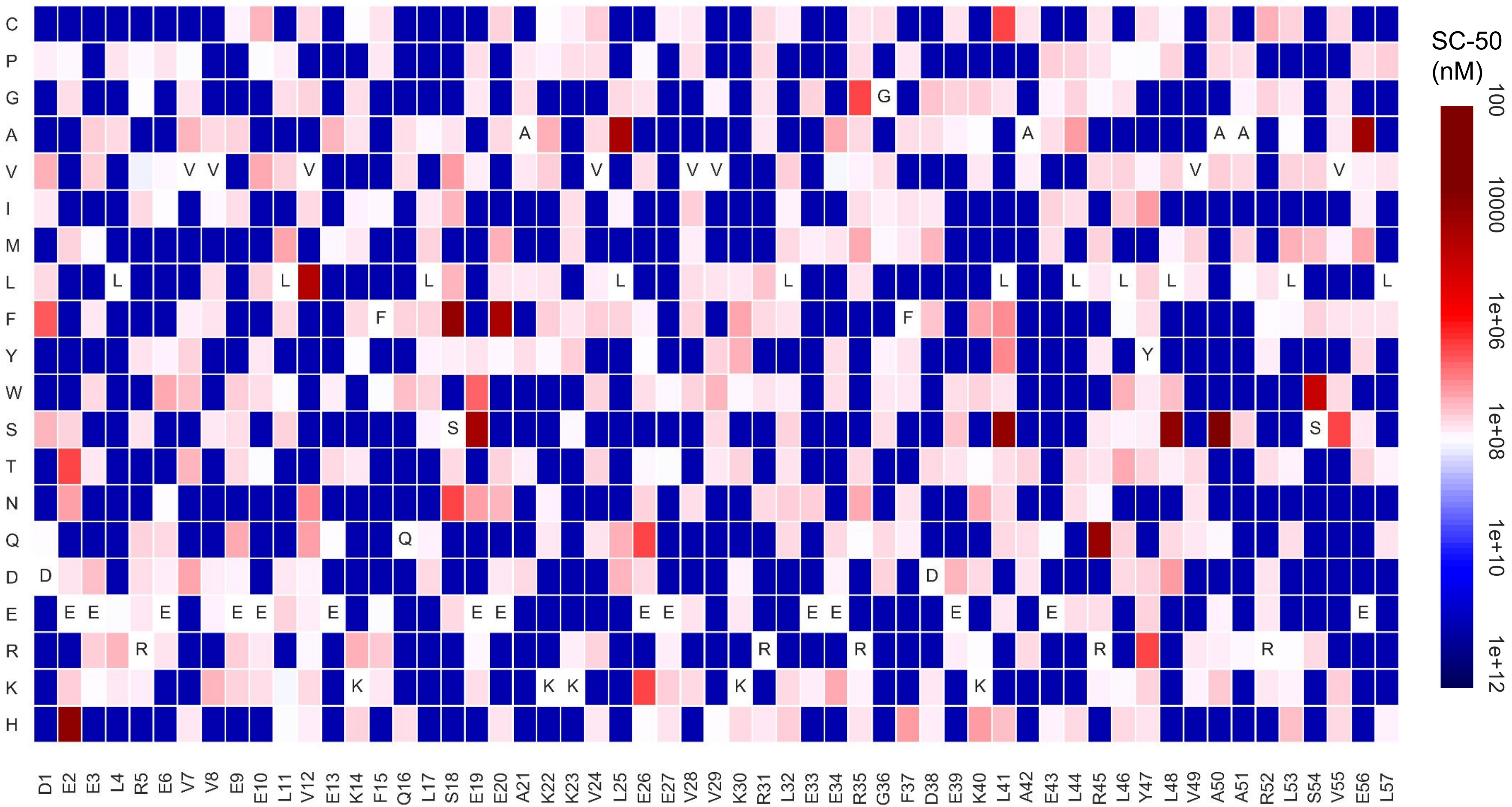

### Minibinder 11

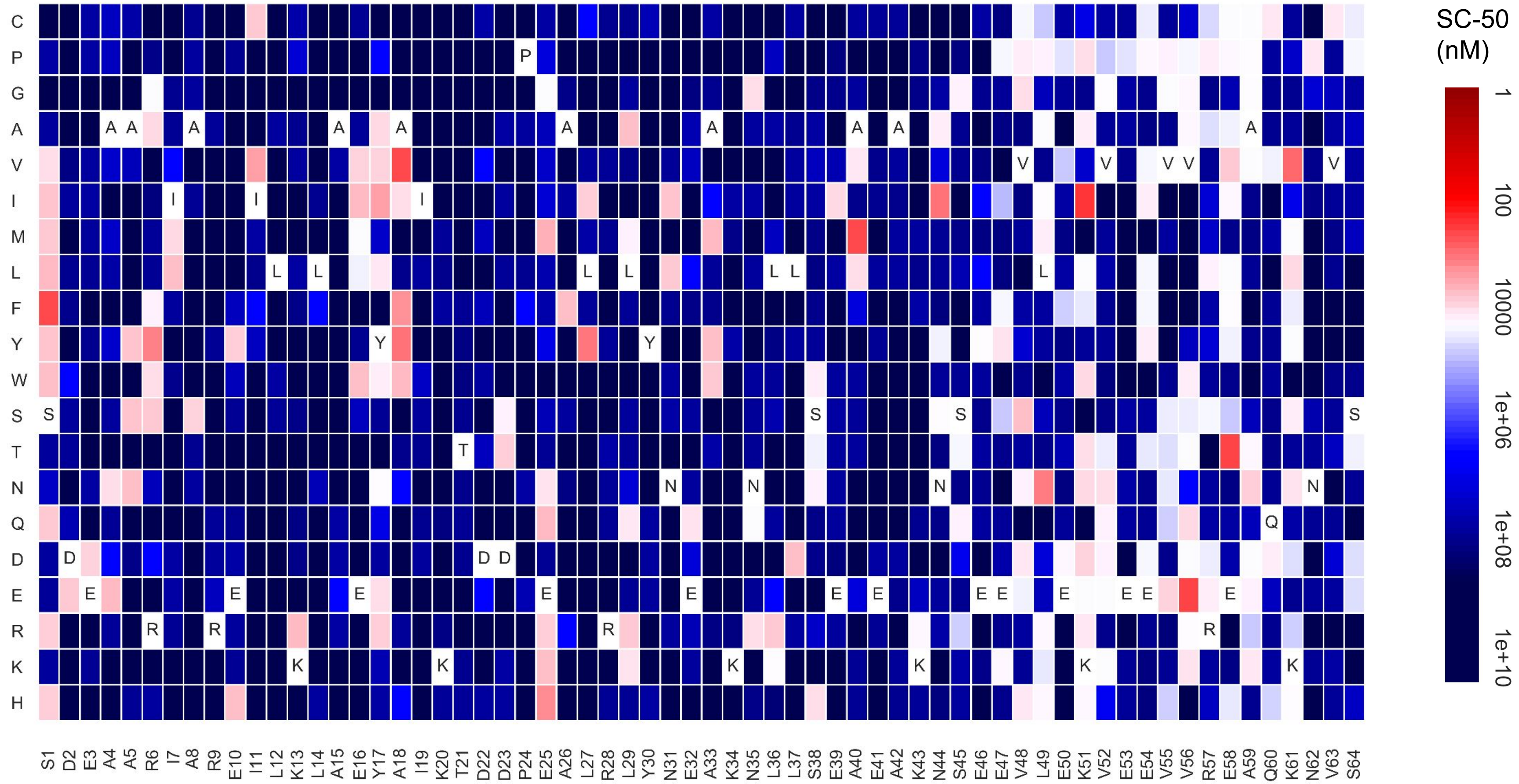

### Full gel from Extended Data Fig. 1

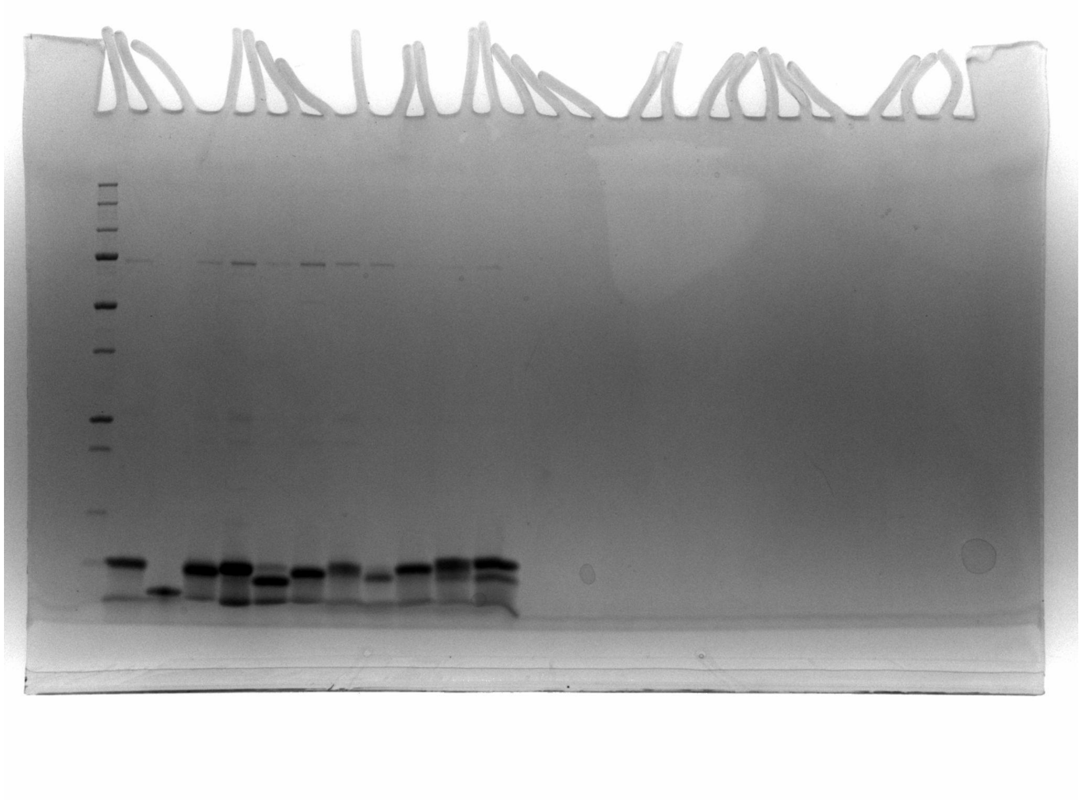

### Full gel from Extended Data Fig. 3

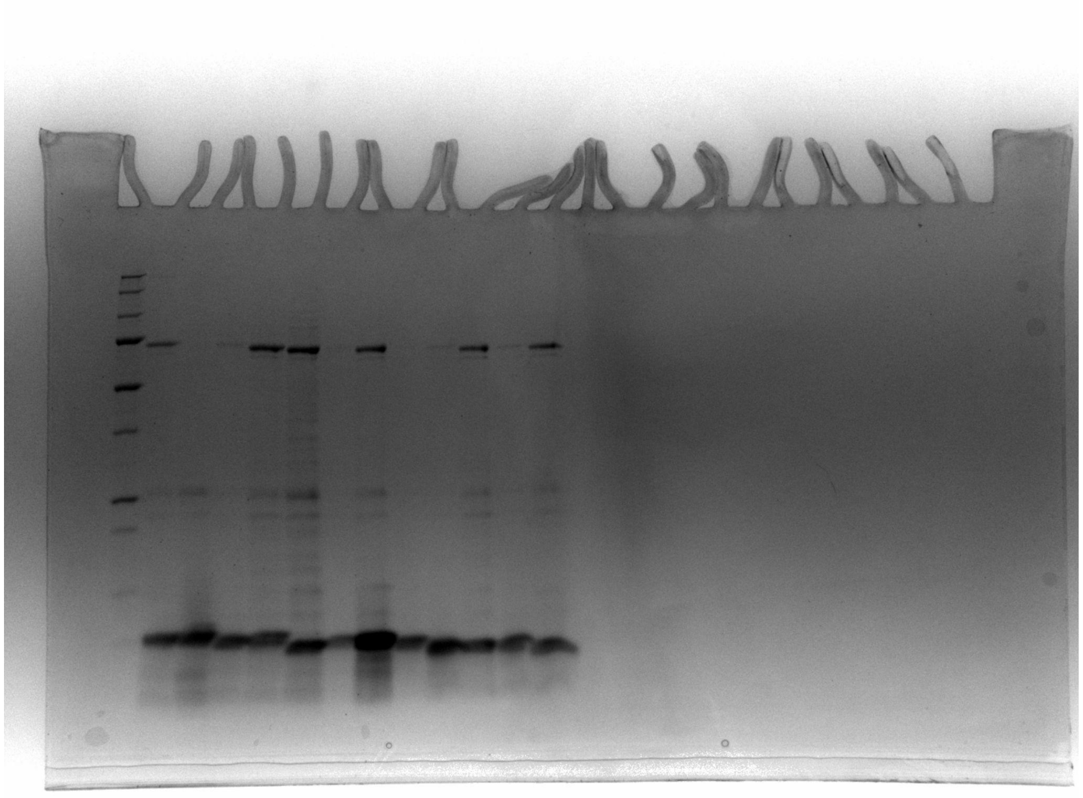

### Full gel from Extended Data Fig. 4

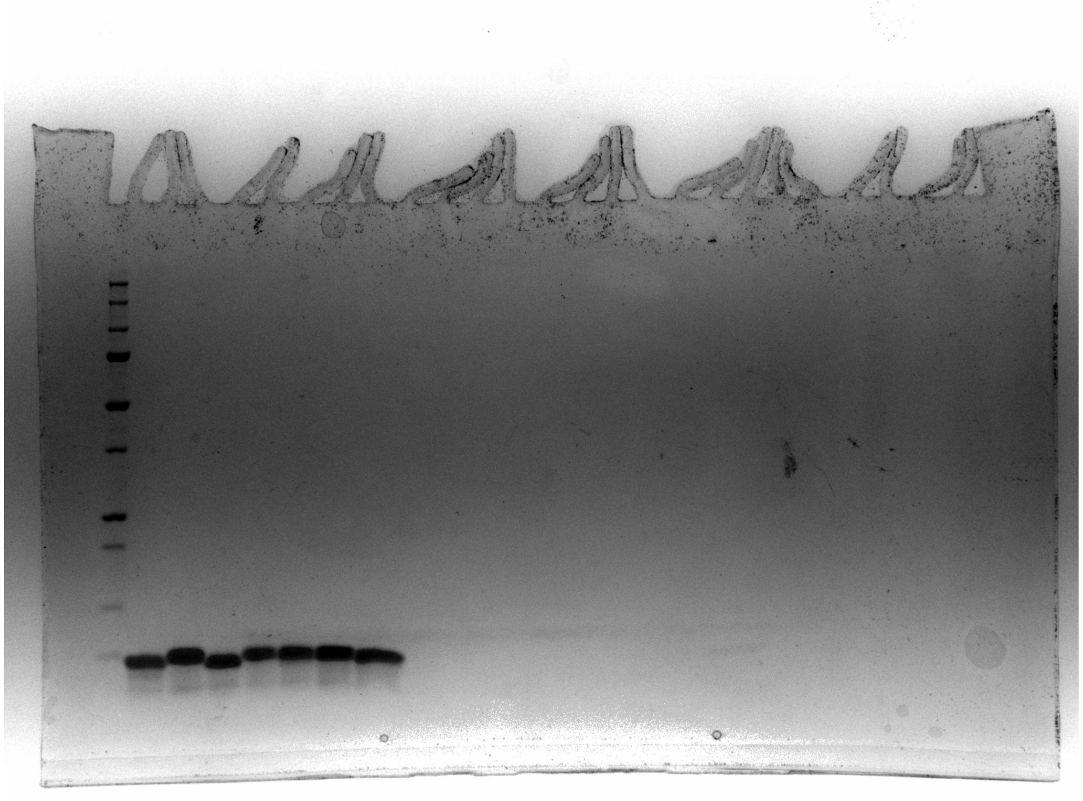
